# Praziquantel activates a schistosome transient receptor potential channel

**DOI:** 10.1101/600791

**Authors:** Sang-Kyu Park, Paul McCusker, Peter I. Dosa, John D. Chan, Jonathan S. Marchant

**Author notes:** Correspondence: Jonathan S. Marchant.

## Abstract

The anthelmintic drug praziquantel (PZQ) is used to treat schistosomiasis, a neglected tropical disease that affects over 200 million people. The target of PZQ in the blood fluke responsible for this disease is unknown. Here, we demonstrate that PZQ activates a transient receptor potential (TRP) channel found in parasitic schistosomes and other PZQ-sensitive parasites.

---

Schistosomiasis (Bilharzia) is a parasitic worm infection that infects millions of people worldwide [1, 2]. Mature parasites living in the vasculature lay eggs which become deposited in host tissues where they trigger local inflammatory responses. Chronic infections become associated with fibrosis and obstructive disease in gastrointestinal tissues and liver (*S. mansoni, S. japonicum*), genitourinary disease (*S. haematobium*), anemia, undernutrition and a heightened risk for many comorbidities. No effective vaccine currently exists for schistosomiasis [3]. The resulting annual disease burden, especially impactful for school-age children, has been estimated a loss of 70 million disability-adjusted life years [1, 2].

In 2017, approximately 100 million people (around 80 million school-aged children) received free preventive treatment for schistosomiasis. The drug praziquantel (PZQ), discovered ∼40 years ago, is the key treatment. PZQ is also used to treat other parasitic infections, caused by tapeworms and trematodes. The clinical formulation of PZQ is a racemate (±PZQ) composed of the enantiomers (*R*)-PZQ and (*S*)-PZQ. (*R*)-PZQ is the anti-schistosomal eutomer, which causes Ca^2+^ influx and spastic paralysis of adult worms, with (*S*)-PZQ regarded as a less active distomer [4, 5]. From a treatment perspective, it is problematic that despite decades of clinical use, as well as demonstration of PZQ resistance in both lab and field, the target of PZQ remains unknown [6]. Resolution of the target of PZQ action in the parasite would be enabling for discovering new anthelmintics and vulnerabilities within signaling pathways engaged by PZQ. This lack of knowledge has proved a longstanding roadblock in this field.

Here, we demonstrate (*R*)-PZQ activates a Ca^2+^ permeable schistosome transient receptor potential (TRP) channel, broadly expressed in PZQ-sensitive flatworms.

## Results

Addition of (*R*)-PZQ (100nM) to adult schistosome worms *in vitro* caused a rapid, spastic paralysis (Figure 1A). Addition of the same concentration of (*S*)-PZQ was ineffective at causing muscle contraction (Figure 1A). These effects reflect the different potency of these enantiomers against schistosomes (EC_50_ for (*R*)-PZQ = 68±7nM, EC_50_ for (*S*)-PZQ= 1.1±0.4µM, ∼16-fold difference, Figure 1B) *in vitro* and *in vivo* [4].

**Figure 1.**
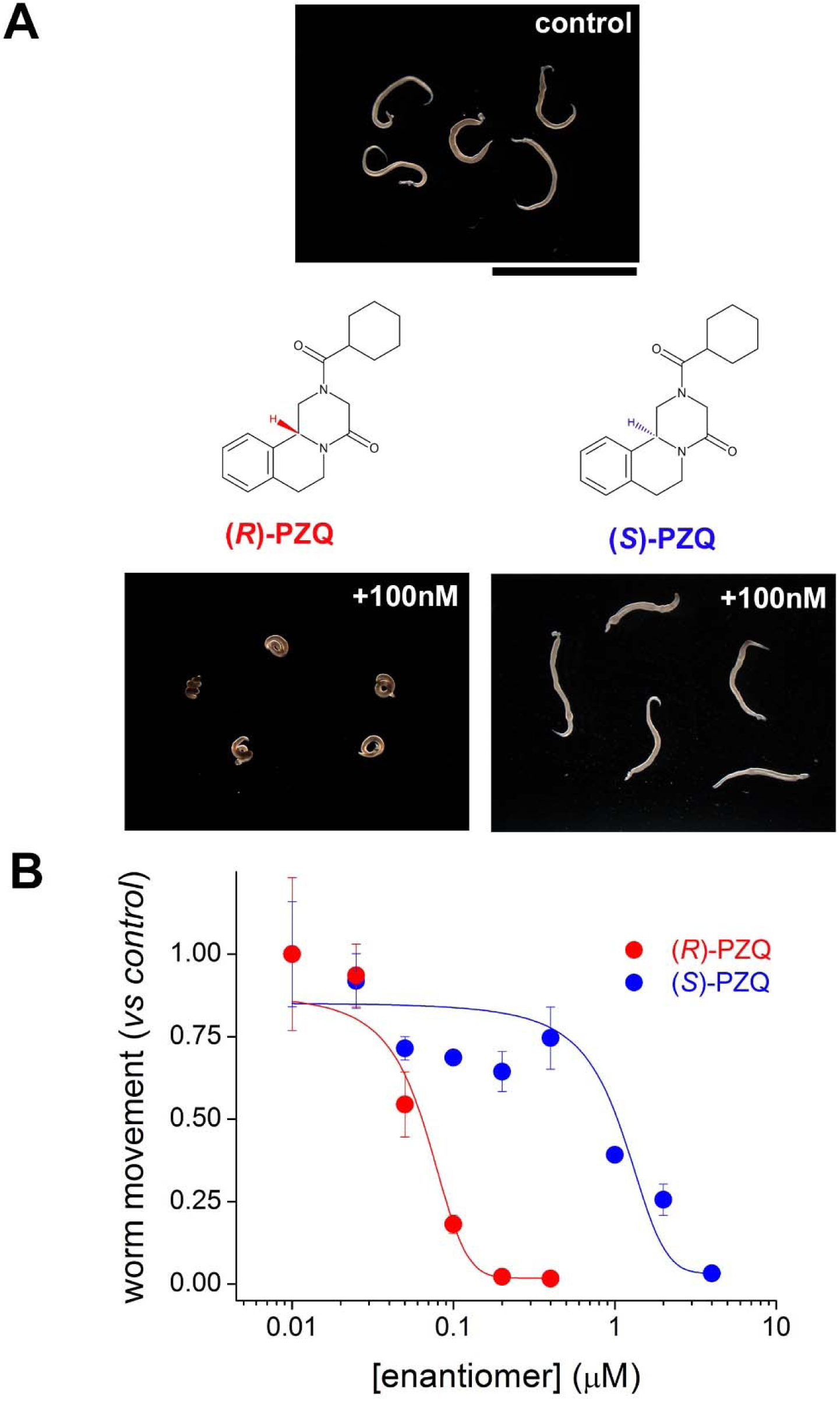
Effects of PZQ enantiomers on schistosome worms *in vitro*. (**A**) Images of five schistosome worms before (top) and after (bottom) addition of a fixed concentration (100 nM) of (*R*)-PZQ or (*S*)-PZQ. Structures of (*R*)-PZQ and (*S*)-PZQ, tetracyclic tetrahydroisoquinolines, highlight chirality. **(B**) Concentration-response relationships for (*R*)-PZQ (red) and (*S*)-PZQ (blue) evoked changes in worm motility.

While no binding site(s) for these enantiomers have been identified in the parasite, there has been considerable recent progress in identifying targets for (*R*)-PZQ and (*S*)-PZQ in the human host [7]. (*R*)-PZQ acts as a partial agonist of the human 5-hydroxytryptamine 2B receptor (5HT_2B_R, [8]) and (*S*)-PZQ is a partial agonist of the human transient receptor potential melastatin-8 channel (hTRPM8, [9]). While regulation of these host targets occurs over a micromolar concentration range (EC_50_ for ±PZQ at 5-HT_2B_R ∼8µM [8], EC_50_ for ±PZQ at hTRPM8 ∼20-25µM [9, 10]), molecular divergence between human and flatworm ligand binding pockets [11, 12] makes it reasonable to anticipate different affinities at homologous schistosome target(s).

Following this logic, we searched for flatworm TRP channels exhibiting homology to hTRPM8, and a BLAST search of the *Schistosoma mansoni* genome prioritized several candidates. These candidates were profiled by transient expression in HEK293 cells using a Ca^2+^ flux assay on a real-time plate reader. HEK293 cells, transiently expressing hTRPM8 (hTRPM8) were used as a positive control for responses to ±PZQ and each enantiomer (Supplementary Figure 1).

A primary screen against a fixed ligand concentration revealed robust responses to ±PZQ with one TRP channel candidate. This candidate, christened *Sm*.TRPM_PZQ_, exhibited robust responses to both ±PZQ and (*R*)-PZQ (Figure 2A) compared with naïve cells (Figure 2B). (*S*)-PZQ also evoked a strong response but with slower kinetics suggestive of stereoselectivity toward the PZQ enantiomers that would be poorly discriminated at the high primary screening concentration (Figure 2A, 50µM). *Sm*.TRPM_PZQ_ did not respond to several known TRP ligands (Figure 2A) including menthol and icilin (human TRPM8 agonists), allyl isothiocyanate (AITC, a TRPA1 agonist) and capsaicin (a TRPV1 activator). Removal of extracellular Ca^2+^, abolished signals, consistent with PZQ-evoked Ca^2+^ entry through *Sm*.TRPM_PZQ_ (Supplementary Figure 2).

**Figure 2.**
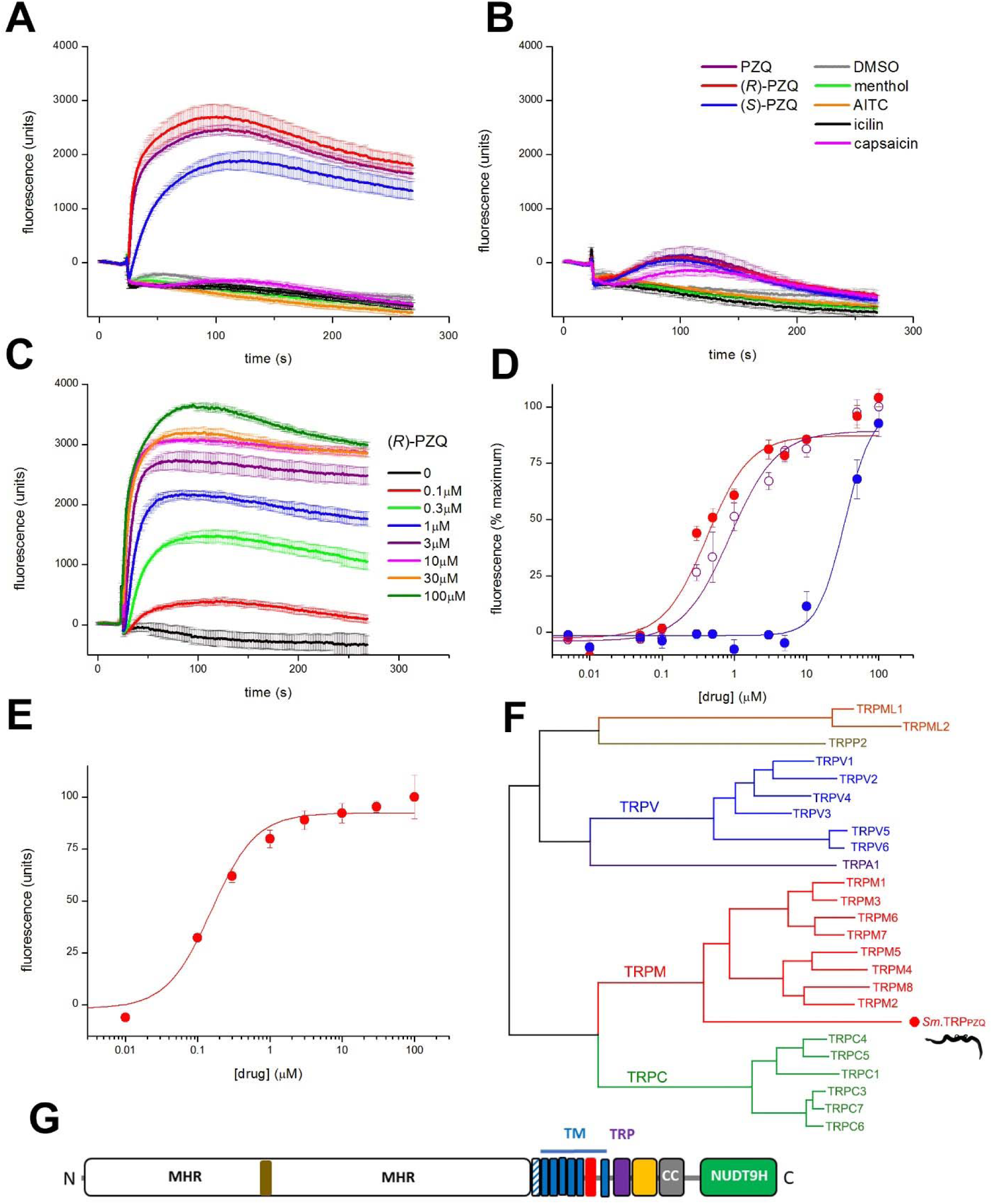
Properties of *Sm*.TRPM_PZQ_. Fluorescence traces in (**A**) HEK293 cells expressing *Sm*.TRPM_PZQ_ (left) and (**B**) untransfected HEK293 cells (right) monitored prior to, and after addition of ±PZQ, (*R*)-PZQ, (*S*)-PZQ and various ligands (all at 50µM, except menthol (500µM), AITC and icilin (100µM)). (**C**) Fluorescence traces showing action of various concentrations (100nM-100µM) of (*R*)-PZQ at *Sm*.TRPM_PZQ_. (**D**) Concentration-response relationships for ±PZQ, (*R*)-PZQ, (*S*)-PZQ at *Sm*.TRPM_PZQ_. (**E**) Concentration-response relationship for (*R*)-PZQ at 37°C. (**F**) Phylogenetic relationship of *Sm*.TRPM_PZQ_ to human TRP channels. Sequence alignments were performed using MUSCLE and phylogeny was inferred using a maximum likelihood method and matrix-based model platform, selecting the iteration with superior log likelihood value. TRP sequences were TRPML1 (Q9GZU1), TRPML2 (Q8IZK6), TRPP2 (Q13563), TRPV1 (Q8NER1), TRPV2 (Q9Y5S1), TRPV3 (Q8NET8), TRPV4 (Q9HBA0), TRPV5 (Q9NQA5), TRPV6 (Q9H1D0), TRPA1 (O75762), TRPM1 (Q7Z4N2), TRPM2 (O94759), TRPM3 (Q9HCF6), TRPM4 (Q8TD43), TRPM5 (Q9NZQ8), TRPM6 (Q9BX84), TRPM7 (Q96QT4), TRPM8 (Q7Z2W7), TRPC1 (P48995), TRPC3 (Q13507), TRPC4 (Q9UBN4), TRPC5 (Q9UL62), TRPC6 (Q9Y210), TRPC7 (Q9HCX4), *Sm*.TRPM_PZQ_ (Smp_246790). (**G**) Domain organization of *Sm*.TRPM_PZQ_. Schematic shows distinct domains identified in recent TRPM2 structures to include: the NH_2_-terminal TRPM homology region (MHR) domain containing an ankyrin-like repeat domain (brown, [16]), the pre S1 helix (shaded), the six transmembrane (TM) spanning helices (S1-S6, blue) with the pore helix between S5 and S6 (red), the TRP domain (purple), the rib and pole helices (yellow), an additional helical domain (CC, black) and the COOH terminal NUDT9H domain.

Full concentration-response curves were performed with (*R*)-PZQ (Figure 2C), (*S*)-PZQ and ±PZQ (Figure 2D). *Sm*.TRPM_PZQ_ was activated by ±PZQ with an EC_50_ of 1.08±0.14µM (Figure 2D) and this activation was stereoselective, with the (*R*)-PZQ evoking Ca^2+^ signals over a considerably lower concentration range (EC_50_ of 597±10nM) than (*S*)-PZQ (EC_50_ of 27.9±3.1µM, Figure 2D). This concentration range, and the clear difference between (*R*)-PZQ and (*S*)-PZQ sensitivity (∼50-fold), is consistent with the action of PZQ on schistosome muscle (Figure 1). Finally, when the incubation temperature was increased to 37°C, (*R*)-PZQ activated *Sm*.TRPM_PZQ_ over a lower concentration range (EC_50_=154±33nM, Figure 2E).

Consistent with our homology search approach, *Sm*.TRPM_PZQ_ displays sequence homology with the hTRPM subfamily (Figure 2F). Sequence analysis revealed an architecture characteristic of TRPM channels (Figure 2G), a family which is well represented within flatworm genomes [13]. Features include a long NH_2_-terminal TRPM homology region (MHR) domain, followed by six predicted transmembrane (TM) domains with a pore-forming re-entry loop between TM5 and TM6, a conserved TRP helix juxtaposed to a coiled-coil region, and a cytoplasmic COOH terminal enzymatic domain (Figure 2F). This enzyme domain shows homology with the human ADP ribose (ADPR) pyrophosphatase NUDT9, a feature characteristic of TRMP2 channels [14-17]. Analysis of *Sm*.TRPM_PZQ_ in transcriptomic datasets evidences expression across various parasite life cycle stages [18], and analysis of flatworm genomes revealed expression of closely related TRPM channels in both free-living and parasitic flatworms that exhibit sensitivity to PZQ (Supplementary Figure 3).

In conclusion, these experiments have identified a Ca^2+^-permeable schistosome TRPM channel activated by (*R*)-PZQ in the nanomolar range.

## Discussion

These data identify a schistosome TRP channel (*Sm*.TRPM_PZQ_) activated by ±PZQ, a discovery that represents the first report of a flatworm target activated by this drug. Identification of a TRPM2-like channel as a ±PZQ target is appealing as it consistent with several known facets of PZQ action.

First, ±PZQ has long been known to cause a sustained Ca^2+^ influx and acute paralysis of schistosome worms [5]. *Sm*.TRPM_PZQ_ is a Ca^2+^ permeable channel that supports long lasting cellular Ca^2+^ signals resulting from Ca^2+^ entry (Figure 2). hTRPM2 is known to exhibit long channel opening times that support substantial Ca^2+^ influx [17, 19]. Second, exposure to ±PZQ causes muscle and tegumental damage in schistosomes [5]. hTRMP2 is a well-known effector of apoptosis being responsive to reactive oxygen species on account of activation by H_2_O_2_ and adenosine 5′-diphosphoribose (ADPR, [17, 20]). Activation of hTRMP2, at the cell surface and within intracellular organelles, causes lysosomal permeabilization and cell death [20-22]. Such actions potentially explain the deleterious effects of ±PZQ on the worm surface. Third, hTRMP2 has a physiological role as a thermosensitive channel sensing non-noxious warmth (reviewed in [19, 23]). Temperature changes are known to depolarize schistosome tegument and muscle [24]. Given the varied environmental transitions experienced during the parasitic life cycle perhaps *Sm*.TRPM_PZQ_ fulfills a critical role in parasite physiology as a thermosensor, which is dysregulated by PZQ. Future work will investigate the mechanisms of endogenous *Sm*.TRPM_PZQ_ channel activation, and synergism between channel ligands and environment. Finally, consistent with *Sm*.TRPM_PZQ_ being a target of this clinically important therapeutic, *Sm*.TRPM_PZQ_ is expressed through various parasite life cycle stages [18] and is present in other flatworms sensitive to PZQ (Supplementary Figure 3).

Identification of *Sm*.TRPM_PZQ_ as a ±PZQ target is significant as this discovery enables execution of target-based screens to discover new scaffolds active at *Sm*.TRPM_PZQ_ that may yield novel anthelmintics. It also provides a molecular handle on key questions for chemotherapy, such as how is *Sm*.TRPM_PZQ_ differentially regulated in juvenile worms which have less sensitivity to PZQ, likely a contributory factor driving drug resistance. A key point is that (*R*)-PZQ acts as a potent activator of *Sm*.TRPM_PZQ_ in the nanomolar range (Figure 2). Existing regulators of hTRPM2 act over the micromolar range - for example, hTRPM2 activates in response to micromolar levels of the endogenous agonist ADPR [17]. This is important as hTRPM2 is an emerging clinical target for several nervous system and inflammatory disorders [17, 19]. Although hTRPM2 does not appear to be activated by PZQ [9], understanding the basis of (*R*)-PZQ affinity for *Sm*.TRPM_PZQ_ and comparing regulation and gating of *Sm*.TRPM_PZQ_ with recently solved TRPM2 structures from vertebrates [14, 15] and other invertebrates [16] may help catalyze drug design at this clinically important target.

Finally, this discovery will prioritize subsequent analyses of other schistosome TRPM channels and splice variants both to assess broader PZQ sensitivity in this clade, and to understand their roles in parasite (patho)physiology which remain largely unexplored.

## Methods

### Reagents

Enantiomers of ±PZQ were resolved following the protocol published by Woelfle *et al.* [25]. All chemical reagents were sourced from Sigma. Cell culture reagents were from Invitrogen. Lipofectamine 2000 was from ThermoFisher.

### Adult schistosome mobility assays

Adult schistosomes were recovered by dissection of the mesenteric vasculature in female Swiss Webster mice previously infected (∼49 days) with *S. mansoni* cercariae (NMRI strain) by the Schistosomiasis Resource Center at the Biomedical Research Institute (Rockville, MD). All animal experiments followed ethical regulations approved by the MCW IACUC committee. Harvested schistosomes were washed in RPMI 1640 supplemented with HEPES (25mM), 5% heat inactivated FBS (Gibco) and penicillin-streptomycin (100 units/mL) and incubated overnight (37°C/5% CO_2_) in vented petri dishes (100×25mm). The following day, movement assays were performed using male worms in six well dishes (∼5 individual worms/3ml media per well). Video recordings were captured using a Zeiss Discovery v20 stereomicroscope coupled to a QiCAM 12-bit cooled color CCD camera controlled by Metamorph imaging software. Recordings (1 minute) of worm motility (4 frames/sec), during addition of various drug concentrations were analyzed as previously described [11]. Data represent mean±standard error for ≥3 independent experiments.

### Cell Culture and transfection

HEK293 cells (ATCC CRL-1573.3) were cultured in DMEM supplemented with 10% fetal bovine serum (FBS), penicillin (100 units/ml), streptomycin (100 μg/ml), and L-glutamine (290 μg/ml). For screening parasite TRP channels, codon-optimized cDNAs (Genscript) were transiently transfected into HEK293 cells using Lipofectamine2000 at a density of 3×10^6^ cells per dish (100mm).

### Ca2+ imaging assays

Ca^2+^ imaging assays were performed using a Fluorescence Imaging Plate Reader (FLIPR^TETRA^, Molecular Devices). Briefly, HEK293 cells (naïve or transfected) were seeded (50,000 cells/well) in a black-walled clear-bottomed poly-d-lysine coated 96-well plate (Corning) in DMEM growth media supplemented with 10% dialyzed FBS. After 24 hrs, growth medium was removed, and cells with loaded with a fluorescent Ca^2+^ indicator (Fluo-4 Direct dye, Invitrogen) by incubation (100 µL per well, 1 h at 37°C) in Hanks’ balanced salt solution (HBSS) assay buffer containing probenecid (2.5 mM) and HEPES (20mM). Drug dilutions were prepared in assay buffer, without probenecid and dye, in V-shape 96-well plates (Greiner Bio-one, Germany). After loading, the Ca^2+^ assay was performed at room temperature. Basal fluorescence was monitored for 20s, then 25μl of each drug was added, and the signal (raw fluorescence units) was monitored over an additional 250s. For quantitative analyses, peak fluorescence in each well was normalized to maximum-fold increase over baseline. Changes in fluorescence amplitude were analyzed using the sigmoidal dose-response function in Origin.

## Acknowledgements

Work was supported by the Marcus Family Foundation, NIH (R21-AI125821, R21-AI130642) and NSF (MCB1615538). Schistosome-infected mice were provided by the NIAID Schistosomiasis Resource Center at the Biomedical Research Institute (Rockville, MD) through NIH-NIAID Contract HHSN272201000005I for distribution via BEI Resources.

