## Supplementary Material for "Praziquantel activates a schistosome transient receptor potential channel"

### Supplementary Information - Park et al.

**Supplementary Figure 1. Activation of human TRPM8 by PZQ enantiomers**. $Ca^{2+}$ responses in HEK293 cells expressing human TRPM8 (hTRPM8) following addition of various concentrations of (**A**) menthol, (**B**) ±PZQ, (**C**) (*R*)-PZQ and (**D**) (*S*)-PZQ.

**Supplementary Figure 2. PZQ-evoked $Ca^{2+}$ signals depend on $Ca^{2+}$ influx.** Responses to PZQ (5μM) in HEK293 cells expressing *Sm*.$TRPM_{PZQ}$ in normal HBSS (purple) and $Ca^{2+}$-free HBSS (black, HBSS supplemented with 1mM EGTA).

**Supplementary Figure 3. *Sm*.$TRPM_{PZQ}$ is present in other PZQ-sensitive flatworms.** Rooted phylogenetic tree showing interrelationship between flatworm TRPM channels clustered with human TRPM channels (hTRPM, dashed line), TRPV1 and TRPA1.Sequences derive from BLAST searches of flatworm genomes and represent the following gene identifiers: Schistosomes [*Schistosoma mansoni* (*Sm*.$TRPM_{PZQ}$ (Smp_246790), Smp_000050, Smp_130890, Smp_333650); *Schistosoma japonicum* (Sjp_0026170); *Schistosoma magrebowiei* (SMRZ_0001736801)], cestodes [*Echinococcus granulosus* (000986600_1_1), *Taenia solium* (000444100_1_1), *Hymenolepis microstoma* (000757100_1_1)], flukes [*Clonorchis sinensis* (csin109609), *Echinostoma caproni* (ECPE_0000226901), *Gyrodactylus salaris* (scf7180006953457)], planarians [*Schmidtea mediterranea* (Smed_v6_17981_0_1), *Dugesia japonica* (Djap_v4_69706)], *Macrostomum_lignano* (Mlig_1509_40127_1, Mlig _1509_54358_1). A subset of other predicted *S. mansoni* TRPM family members related to *Sm*.$TRPM_{PZQ}$ are shown in purple.

**Supplementary Figure 1**

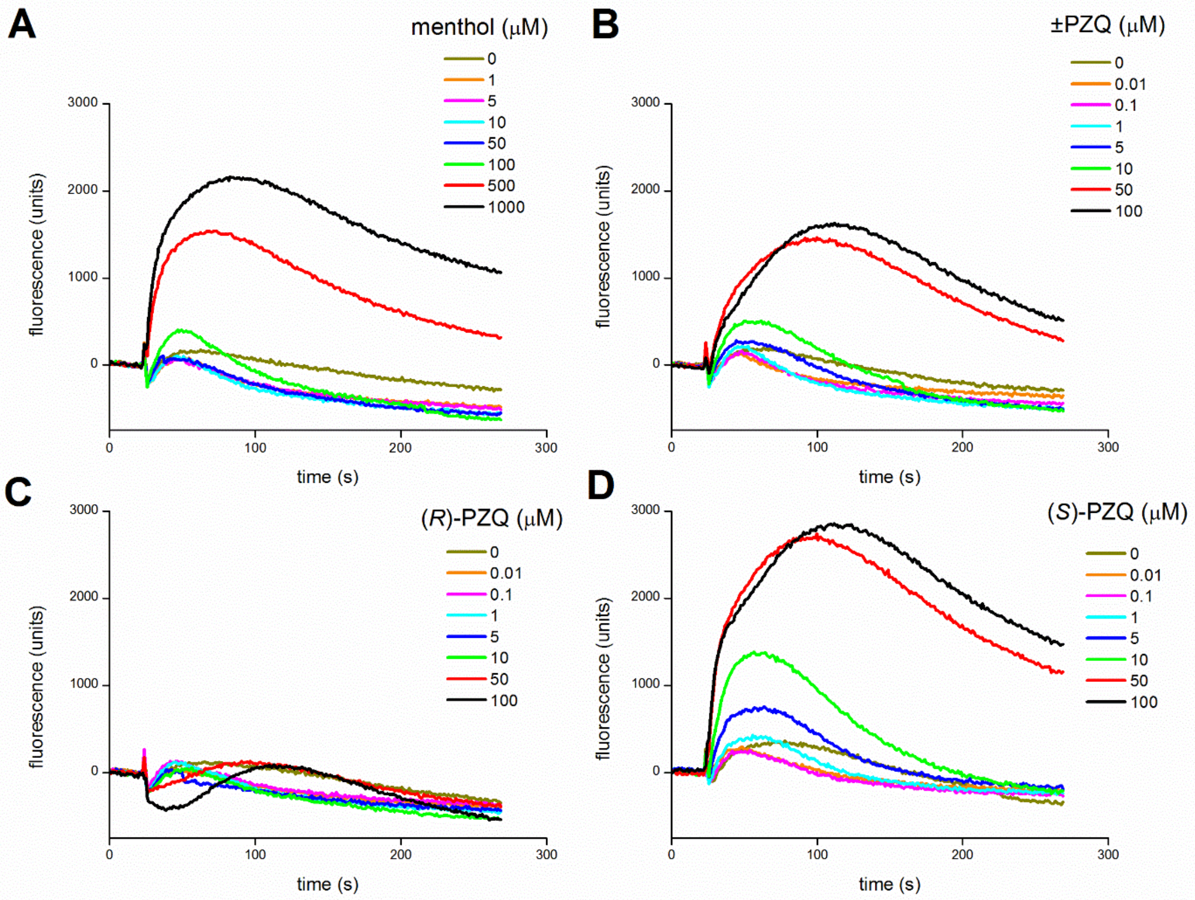

**Supplementary Figure 2**

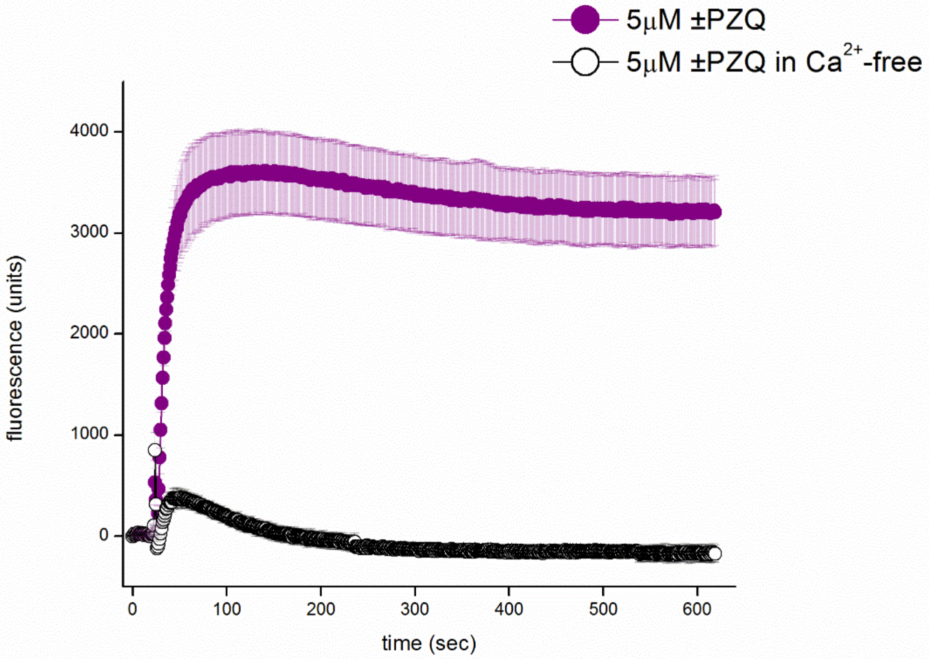

**Supplementary Figure 3**

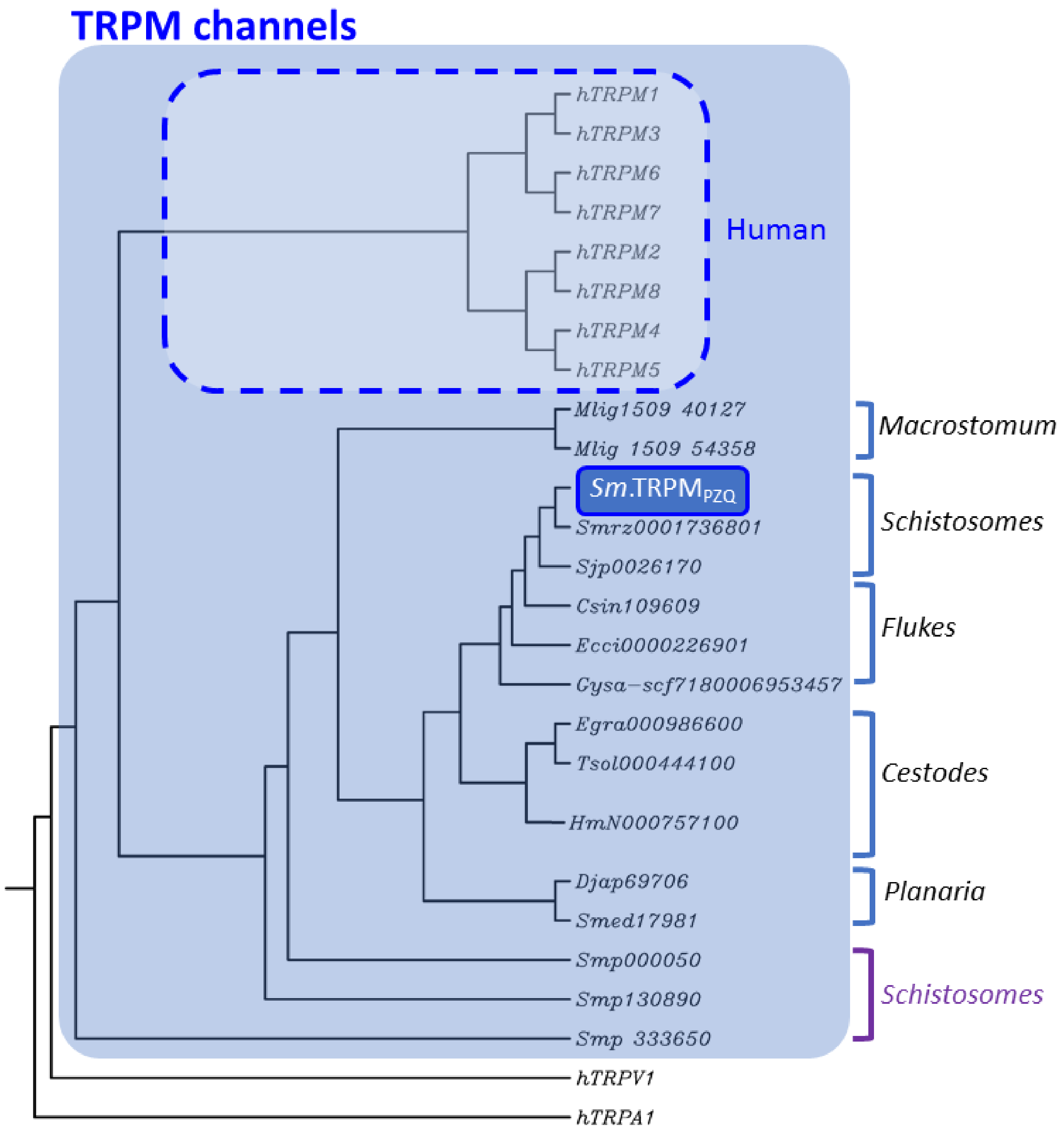
